# Enhanced nitrous oxide production in denitrifying *Dechloromonas aromatica* strain RCB under salt and alkaline stress conditions

**DOI:** 10.1101/497941

**Authors:** Heejoo Han, Bongkeun Song, Min Joon Song, Sukhwan Yoon

## Abstract

Salinity and pH are important environmental parameters with direct and indirect impacts on the viability and metabolic activities of microorganisms. In this study, the effects of salt and alkaline stresses on the kinetic balance between nitrous oxide (N_2_O) production and consumption in the denitrification pathway of *Dechloromonas aromatica* strain RCB were examined. N_2_O accumulated transiently only in insignificant amounts at low salinity (≤0.5% NaCl) and circumneutral pH (7.0 and 7.5). Incubation at 0.7% salinity resulted in substantially longer lag phase and slower growth rate, along with the increase in the amounts of transiently accumulated N_2_O (15.8±2.8 μmoles N_2_O-N/vessel). Incubation at pH 8.0 severely inhibited growth and resulted in permanent accumulation of 29.9±1.3 μmoles N_2_O-N/vessel from reduction of 151±20 μmoles NO_3_^-^. The transcription analyses observed decreased *nosZ*/(*nirS*_1_+*nirS*_2_) ratios coinciding with N_2_O accumulation. The N_2_O consumption rates of resting *D. aromatica* cells subjected to the salt and alkaline stress conditions were significantly lower than the rates of N_2_O production from NO_2_^-^ reduction at N_2_O / NO_2_^-^ concentration of 0.1 mM, but not at a higher concentration (1.0 mM). These results indicate that alteration in N_2_O consumption kinetics was another cause of enhanced N_2_O production observed under the stress conditions. The findings in this study suggest that canonical denitrifiers may become a significant N_2_O source when faced with abrupt environmental changes.

**Importance:** Denitrification is one of the two most significant biological sources of N_2_O. In a complete denitrification pathway, N_2_O generated from stepwise reduction of NO_3_^-^ via NO_2_^-^ and NO is simultaneously reduced to N_2_. Thus, N_2_O evolution is often kept minimal in cultivation of canonical denitrifiers at their optimal growth conditions. This study discovered that this orchestration of the reactions in the denitrification cascade is disrupted in *Dechloromonas aromatica*, an environmentally abundant complete denitrifier, as a stress response to elevated salinity and pH impairing cell viability and growth. This disruption in the kinetic balance resulted in >100-fold increase in the amounts of N_2_O transiently or permanently produced during the course of denitrification. As rapid fluctuations of salinity and pH are often observed in both natural ecosystems and engineered systems, N_2_O emissions from the canonical denitrifiers responding to environmental stresses may be a significant, but previously unrecognized, source of N_2_O.

## Introduction

Nitrous oxide (N_2_O), currently constituting 320 ppbv of the atmosphere, is a potent greenhouse gas with approximately 300 times higher global warming potential than carbon dioxide and the most influential ozone depletion agent (1, 2). Due to the ever increasing global demand for food production, which inevitably accompanies increasing demands for nitrogen fertilizers, a steady increase in anthropogenic N_2_O emission is anticipated in the foreseeable future (3). The occurrences of summer heat waves in the Arctic regions that have become increasingly regular in the recent years may also contribute to substantial increase in N_2_O emissions from thawing permafrost tundra soils, providing positive feedback to global warming (4). Therefore, continued research efforts are warranted for identification of yet unknown N_2_O sources and development of environmental management strategies to minimize N_2_O emissions from its sources.

Unlike carbon dioxide, N_2_O emitted to the atmosphere is predominantly of biological origin (5). Many different reaction pathways of the biological nitrogen cycle are responsible for production of N_2_O to some degree; of these pathways, denitrification and nitrification are indisputably the two most important contributors (6). The nitrification pathway consists of two separate reactions, ammonia (NH_3_) oxidation to nitrite (NO_2_^-^) and NO_2_^-^ oxidation to nitrate (NO_3_^-^), which had been known to take places in separate groups of organisms until the recent discovery of the comammox (complete ammonia oxidizing bacteria) *Nitrospira* spp. (7, 8). Multiple N_2_O production mechanisms have been identified in the nitrification pathway, including detoxification of nitric oxide (NO) produced from hydroxylamine (NH_2_OH) oxidation and nitrifier denitrification (9, 10). In the denitrification pathway, the stepwise reduction NO_3_” and NO_2_^-^ to dinitrogen (N_2_) via NO and N_2_O, N_2_O may be produced as a stable intermediate or the final product (6, 11–13). A canonical denitrifier is capable of catalyzing each reduction step in the pathway using a complete suite of the denitrification enzymes; however, partial denitrifiers, lacking nitrous oxide reductase gene, are not rare (11, 14). The partial denitrifiers, including fungal denitrifiers, produce N_2_O as the final product of denitrification (12, 15). Even in the canonical denitrifiers, trace amounts of N_2_O is often transiently detected during the course of denitrification and certain environmental conditions, e.g., acidic pH and low copper bioavailability, may amplify N_2_O production (13, 16–18). Although the relative contributions of nitrification and denitrification to N_2_O emissions from the natural (e.g., wetland soils and freshwater sediments) and anthropogenic sources (e.g., agricultural soils and wastewater treatment plants) are still under controversy, denitrification is generally perceived to be more important N_2_O-emitting pathway in environments under permanent or periodic anoxia (19–21).

The disruption of the N_2_O balance and subsequent increase in transient / permanent N_2_O production at acidic pH was repeatedly observed in pure denitrifying cultures and environmental samples and enrichments (22–25). the effect of other environmental stresses, e.g., alkaline pH and salinity, on N_2_O release from denitrification of a canonical denitrifier has not yet been thoroughly investigated. Periodic fluctuation or abrupt changes of environmental parameters, e.g., pH and salinity, are not unexpected in natural and built environments where denitrification occur. In activated-sludge WWTPs, fluctuations in pH within circumneutral range (6.0-8.0) are often recorded (26). Abrupt increases in soil salinity due to intrusions of seawater into fertilized coastal agricultural areas have recently become increasingly frequent as consequences of climate change (27). Although impacts of such temporal shifts in the environmental conditions on the microbial compositions have been observed to be substantial in various ecological niches, little has been investigated regarding the ecophysiological responses of isolates representing key functional groups of biogeochemical cycling to such stresses (28–32).

*Dechloromonas* spp. and closely related taxa affiliated to the *Betaproteobacteria* class are often among the dominant denitrifying population in activated sludge tanks of wastewater treatment plants and are also found in abundance in soils and freshwater sediments (33–35). The isolates affiliated to this organismal group, when monitored for progression of denitrification at their optimal growth conditions (i.e., neutral pH, optimal temperature, and low salinity) in batch experiments, exhibited minimal transient N_2_O accumulation, presumably owing to their high affinity to N_2_O (17, 36–38). Perhaps, to maximize the energy yield from limited amounts of NO_3_^-^/NO_2_^-^, these denitrifiers may have evolved to maintain kinetic balance between production and consumption of N_2_O, such that loss of this energy-yielding terminal electron acceptor is minimized (39).

In this study, we have investigated N_2_O production from the denitrification pathway in *D. aromatica* strain RCB, a model organism representing the *Dechloromonas-like* denitrifiers, upon incubation under salt and alkaline stresses. When strain RCB was exposed to the salt and alkaline stresses significantly inhibiting cell growth, N_2_O evolution substantially increased due to the disrupted balance between the N_2_O-producing and reducing steps of denitrification. Both the transcription analyses using reverse transcription polymerase chain reaction (RT-qPCR) and the measurements of the reaction kinetics at the stress conditions explained disruption of the N_2_O balance in strain RCB subjected to growth-limiting environmental stresses. This previously underappreciated N_2_O production mechanism may account for substantial portions of N_2_O emissions from the environments with fluctuation pH and/or salinity.

## Results

### Effect of salt and alkaline stress on N_2_O evolution from denitrifying *D. aromatica* strain RCB

The effects of salt and alkaline stresses on growth and N_2_O production of *D. aromatica* strain RCB were monitored in anaerobic batch cultures with 10 mM acetate and 5.0 mM NO_3_^-^ provided as the sole electron donor and acceptor, respectively (Table 1, Fig. 1, Fig. S1). No significant accumulation of N_2_O was observed in the cultures incubated with salt (NaCl) concentrations up to 5.0 g L^-1^ despite the lower observed growth rates. At the NaCl concentration of 7.0 g L^-1^, a significant increase (*p*<0.05) in N_2_O accumulation associated with severe growth inhibition (the lag phase prolonged to ~222 hours and growth rate diminished to 0.032 hr^-1^) was observed. The accumulated N_2_O was eventually reduced completely, presumably to N_2_; however, the maximum amount of accumulated N_2_O reached 15.8±2.8 μmoles N_2_O-N/vessel at the peak (t = 222 h). No exponential growth took place at 9.0% NaCl, although 1.97±0.38 μmoles N_2_O-N/vessel (0.40% of added NO_3_^-^) was produced and remained in the medium until the end of incubation. Although neither NO3^-^ nor NO2^-^ concentration was monitored in these preliminary experiments, the absence of cell growth suggested that only insignificant portion of initially added NO_3_^-^ was reduced. The elevated pH also had unexpectedly pronounced impacts on N_2_O production and growth of *D. aromatica* strain RCB. At pH 8.0, the cell density reached only up to an OD_600nm_ value of 0.013 (from an initial value of 0.001) and exponential growth was not observed. The amounts of the accumulated N_2_O-N in the vessels reached a maximum of 31.7±2.06 μmoles and the produced N_2_O permanently remained in the vessels until the ends of the experiments.

**Fig. 1:**
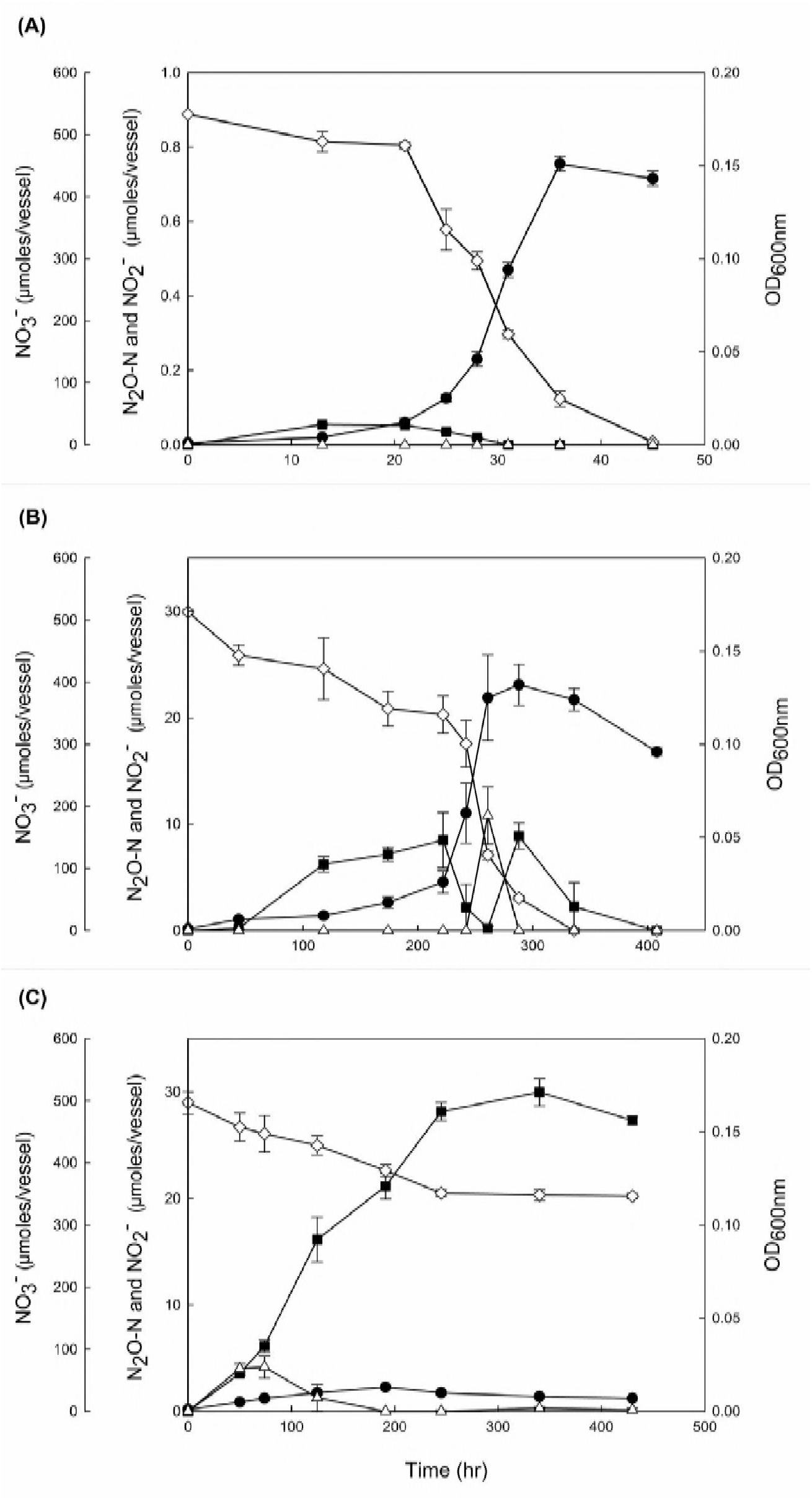
Reduction of 5 mM NO_3_^-^ (in 100 mL liquid medium in 160 mL vessel) by the denitrifying *Dechloromonas aromatica* RCB cultures incubated at **(A)** the control condition (0.05% w/v NaCl, pH 7.0), **(B)** the salt stress condition (0.70% w/v NaCl, pH 7.0) and (C) the alkaline stress condition (0.05% w/v NaCl, pH 8.0). The amounts of NO_3_^-^ (◊), NO_2_^-^ (△) and N_2_O-N (■) and the absorbances at 600 nm (•) were monitored until the nitrogen oxides were depleted or upon confirmation of the termination of denitrification reaction. The data points represent the averages of triplicate cultures and the error bars the standard deviations of the triplicate measurements.

**Table 1.** N_2_O-N accumulation and cell growth in denitrifying *D. aromatica* strain RCB cultures grown on 10 mM acetate and 5 mM NO_3_^-^ at varying NaCl concentrations and pH

| NaCl concentration (g L<sup>-1</sup>) | pH | Maximum N<sub>2</sub>O-N accumulation (μmoles/vessel) | Duration of Incubation (h) | Residual N<sub>2</sub>O (μmoles/vessel) | Maximum OD<sub>600nm</sub> | Specific growth rate (h<sup>-1</sup>) |
| --- | --- | --- | --- | --- | --- | --- |
| 0.5 | 7.0 | 0.05 (0.24)<sup>a</sup> | 51 | 0 | 0.165 (0.010) | 0.226 (0.008) |
| 3.0 | 7.0 | BDL<sup>b</sup> | 46 | 0 | 0.190 (0.004) | 0.177 (0.012) |
| 5.0 | 7.0 | BDL | 50 | 0 | 0.177 (0.006) | 0.131 (0.010) |
| 7.0 | 7.0 | 15.8 (2.83) | 334 | 0 | 0.141 (0.004) | 0.032 (0.001) |
| 9.0 | 7.0 | 2.08 (0.70) | 407 | 1.97 (0.38) | 0.010 (0.000) | NA<sup>c</sup> |
| 0.5 | 7.5 | 0.05 (0.00) | 46 | 0 | 0.163 (0.006) | 0.135 (0.012) |
| 0.5 | 8.0 | 31.7 (2.06) | 424 | 31.6 (2.07) | 0.018 (0.000) | NA |

<sup>a</sup>The numbers in the parentheses are standard deviations of triplicate cultures.
<sup>b</sup>Below detection limit.
<sup>c</sup>The exponential growth was not observed.

Based on these results, a salt stress condition (7.0 g L^-1^) and an alkaline stress condition (pH 8.0) were selected for closer examination, whereby the cell growth and progression of denitrification reaction were monitored in *D. aromatica* strain RCB cultures (Fig. 1). In the control cultures (0.5 g L^-1^ of NaCl, pH 7.0), NO_3_^-^ and its intermediate, NO_2_^-^ and N_2_O, were completely reduced within 45 hours. Accumulation of N_2_O occurred during the early phase of growth (t<31 h), reaching the maximum of 0.054±0.002 μmoles N_2_O-N/bottle at t = 13 h; however, this amount was merely ~0.001% of the total amount of NO_3_^-^ reduced beyond NO_2_^-^. The concentration of NO_2_^-^ remained below the detection limit throughout the incubation.

In the batch cultures of *D. aromatica* strain RCB incubated at the salt stress condition, the lag phase was remarkably prolonged and the exponential growth did not take off until >200 h after inoculation. The exponential growth rate, 0.032±0.001 h^-1^, was also significantly lower than that of the control, 0.218±0.014 h^-1^ (*p*<0.05). Substantial accumulation of N_2_O (up to 8.5±2.6 μmoles N_2_O-N/bottle) was observed during the lag phase (44 – 222 h), but accumulated N_2_O was almost completely consumed (to 0.19 ± 0.018 h^-1^) during the belated exponential growth phase. Unexpected N_2_O accumulation was observed in the early stationary phase (up to 8.9 ± 2.6 μmoles N_2_O-N/bottle) before N_2_O was eventually consumed after depletion of NO_3_^-^. A modest transient accumulation of NO_2_^-^ (up to 10.8 ± 2.7 μmoles/bottle) observed towards the end of the exponential growth phase (t=261 h) suggested a temporal imbalance between NO_3_^-^-to-NO_2_^-^ reduction and NO_2_^-^ reduction.

The growth of *D. aromatica strain* RCB was substantially inhibited at the alkaline stress condition (pH 8.0). Despite the absence of discernable exponential growth phase and the low cell density that was limited to a maximum of OD_600nm_= 0.013±0.001, NO_3_^-^ concentration decreased from 497.1±1.0 to 346.6±4.0 μmoles NO_3_^-^/bottle over 250 h of incubation. Accumulation of NO_2_^-^ remained modest (<4.16±1.0 μmoles NO_2_^-^/bottle and was sustained only at the initial phase of incubation (<191 h). A significantly larger fraction of reduced NO_3_^-^ (18.7±2.4%) was recovered as N_2_O at the end of incubation than any other conditions examined; however, the reaction stoichiometry suggested against the possibility that the nitrous oxide reductase was completely inactivated. In contrast to the other incubation conditions, denitrification activity and cell growth were terminated after 245 h as indicated by the stable NO_3_^-^ and N_2_O concentrations and decreasing cell density, presumably due to the modest increase in the pH to ~8.2 (data not shown).

### RT-qPCR analyses of *nirS, norB*, and *nosZ* transcription

Reverse transcription quantitative polymerase chain reaction (RT-qPCR) analyses were performed to examine whether the increased transient / permanent accumulation of N_2_O at the stress conditions can be explained in terms of transcriptional regulation. The *nirS*_1_, *nirS*_2_, *norB* and *nosZ* transcripts were quantified with the samples collected from the control, salt-stressed, and alkaline-stressed cultures at different stages of growth (Fig. 2). The two copies of *nirS* genes, with 59% translated amino acid identity, exhibited distinct transcriptional patterns (14). Transcription of *nirS*_1_ appeared to be inducible by nitrogen oxides, whereas transcription of *nirS*_2_ was maintained relatively constant throughout the incubation at all three experimental conditions. In the samples taken during the exponential growth phases (t = 31 h for the control cultures and t = 288 h for the salt-stress cultures), the *nirS*_1_ transcription levels were at least 4.8-fold higher than those of *nirS*_2_. Both *nirS*_1_ and *nirS*_2_ were relatively highly expressed throughout the incubation at pH 8.0 despite severely inhibited cell growth (Fig. 2).

**Fig. 2:**
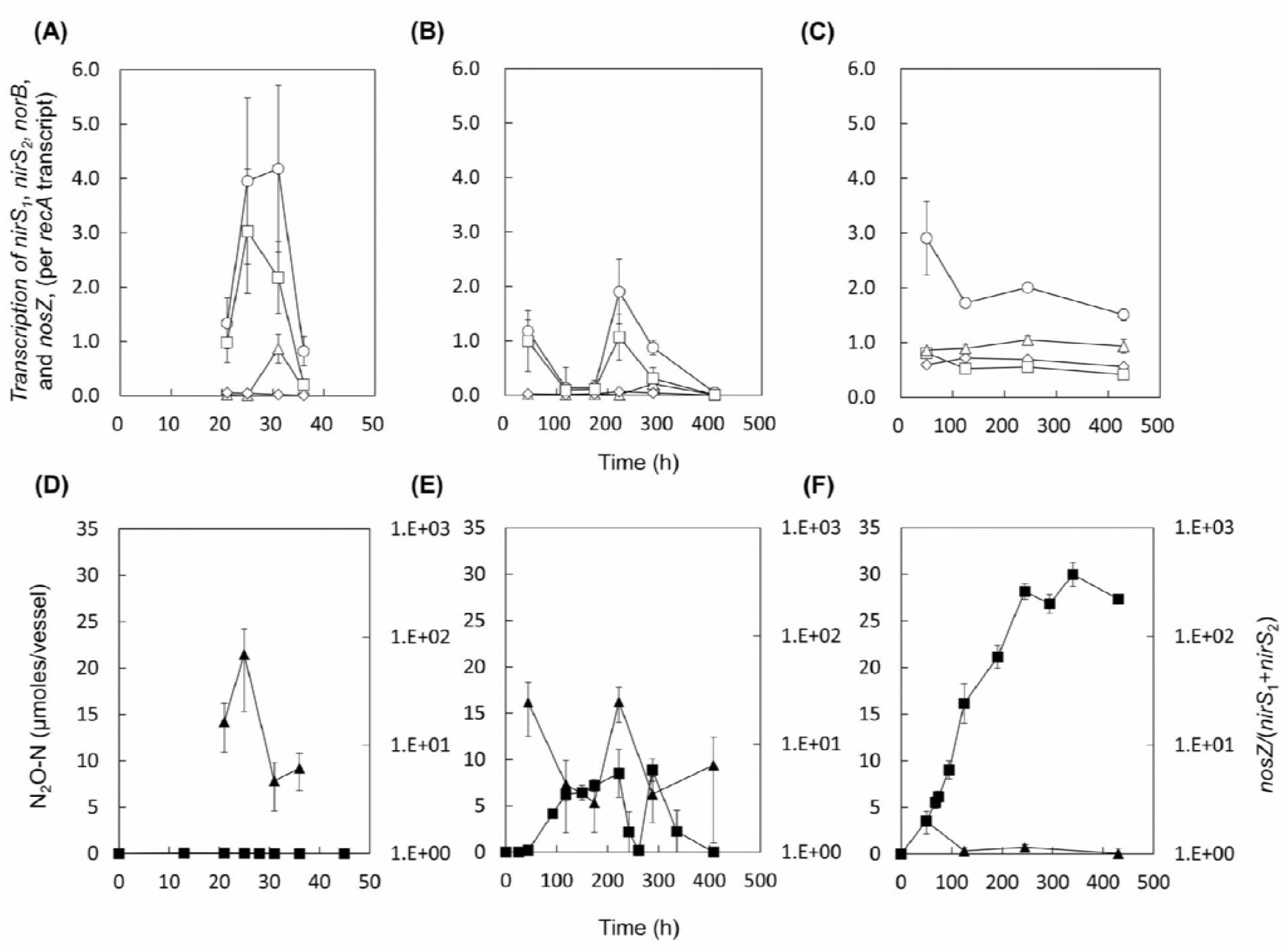
Transcription profiles of *nirS_1_* (△), *nirS_2_* (◇), *norB* (◻), and *nosZ* (○) in the *D. aromatica* strain RCB cultures incubated at **(A)** the control condition **(B)** the salt stress condition and **(C)** the alkaline stress condtions. The samples for the RT-qPCR analyses were withdrawn from the same culture vessels used to construct Figure 1. The transcript copy numbers of the denitrification genes were normalized with the copy numbers of *recA* transcripts. The data points represent the averages of three biological replicates processed independently through RNA extraction and reverse transcription procedures. The error bars represent the standard deviations of the triplicate samples. The values of *nosZ* / (*nirS*_1_+*nirS*_2_) (▲) calculated with the *nirS*_1_, *nirS*_2_, and *nosZ* transcription data in **(A), (B)**, and **(C)** are presented in **(D), (E)**, and **(F)**, respectively. The errors were calculated using the propagation of error. The amounts of N_2_O (◼) are included for convenient comparison between the datasets.

At all experimental conditions, *nosZ* was always the most highly transcribed of the denitrification genes. Similar to the transcriptional pattern of *nirS*_1_, *nosZ* transcription fluctuated during the course of incubation, with increased transcription observed during the periods of active denitrification (characterized by NO_3_^-^ reduction and/or N_2_O production) and down-regulated transcription upon depletion of the nitrogen oxides. In the salt stress samples, the period of down-regulated *nosZ* transcription (t = 118-174 h) during the lag phase coincided with the period of N_2_O accumulation. The peak *nosZ*/(*nirS*_1_+ *nirS*_2_) value was observed at the onset of the exponential growth phase, consistent with the net N_2_O consumption observed between 177-222 h. The *nosZ*/(*nirS*_1_+ *nirS*_2_) values were consistently sustained relatively low in the alkaline stress sample throughout the incubation, agreeing with persistent N_2_O accumulation observed at this incubation condition. The transcription of *norB* generally followed the same trends as the transcription of *nosZ*, suggesting that the expressions of these two genes may be co-regulated in strain RCB; however, *norB* transcription did not exhibit any evident relation with N_2_O production. Although the periods with low *nosZ*/(*nirS*_1_+ *nirS*_2_) values generally agreed with the periods of the net N_2_O productions, the result from the transcription analysis can only serve as circumstantial evidence.

### NO_2_^-^ and N_2_O reduction kinetics of resting *D. aromatica* strain RCB cells under imposed salt and alkaline stress conditions

A set of resting cell kinetic experiments were performed to examine the effects of high salt concentration and alkaline pH on the activities of N_2_O-producing and reducing reactions mediated by *D. aromatica* strain RCB (Table 2). The rates of N_2_O production from NO_2_^-^ reduction were statistically indistinct across the resting cells with or without salt or pH amendments (one-way ANOVA, p>0.05). The substrate concentration did not have significant effect on the N_2_O production rates, suggesting that the whole-cell half-saturation constant (K_m,app_) for NO_2_^-^ reduction is substantially lower than 0.1 mM at all examined conditions. Contrastingly, the rates of N_2_O consumption were significantly lower at both stress conditions as compared to the control condition (*p*<0.05). The potential N_2_O consumption rates measured with the control cultures were statistically indistinct at the two different dissolved N_2_O concentrations (1.0 mM and 0.1 mM) (*p*>0.05). These rates were significantly higher than the measured N_2_O production rates (*p*<0.05), supporting the near absence of transient N_2_O accumulation in the control cultures. In both salt- and pH-stressed cultures, the N_2_O consumption rates were significantly lower for the cultures amended with 0.1 mM N_2_O than those amended with 1.0 mM N_2_O (*p*<0.05). While the N_2_O reduction rates were still higher than the N_2_O production rates at 1.0 mM, the reduction rates at 0.1 mM were significantly lower than the N_2_O production rates (*p*<0.05). Although the Michaelis-Menten kinetics parameters were not determined, these observations suggest that the affinity of the N_2_O reductase in *D. aromatica* strain RCB was substantially altered by both salt and alkaline stresses. As the concentration of N_2_O was sustained below 0.3 mM throughout the entire incubation periods at all three incubation conditions (Fig. 1), the alteration of the N_2_O consumption kinetics in the salt- and pH-amended cultures provides additional, more succinct, mechanistic explanation for observed N_2_O accumulation.

**Table 2.** N_2_O production and reduction rates of resting *D. aromatica* strain RCB cells amended with NaCl (to 0.7% w/v) or NaOH (to pH 8.0).

| Substrate concentration<br>(mM NO<sub>2</sub><sup>-</sup><br>or mM N<sub>2</sub>O-N) | Amendment | N<sub>2</sub>O production rate<br>from NO<sub>2</sub><sup>-</sup> reduction<br>(μmoles N<sub>2</sub>O min<sup>-1</sup><br>mg protein<sup>-1</sup>) | N<sub>2</sub>O consumption rate<br>(μmoles min<sup>-1</sup><br>mg protein<sup>-1</sup>) |
| --- | --- | --- | --- |
| 0.1 | Not amended | 0.52 (0.02)<sup>a</sup> | 0.84 (0.07) |
|  | 7.0 g L<sup>-1</sup> NaCl | 0.46 (0.05) | 0.27 (0.02) |
|  | pH 8.0 | 0.51 (0.04) | 0.31 (0.03) |
| 1.0 | Not amended | 0.55 (0.05) | 0.91 (0.06) |
|  | 7.0 g L<sup>-1</sup> NaCl | 0.52 (0.05) | 0.57 (0.02) |
|  | pH 8.0 | 0.45 (0.05) | 0.66 (0.02) |

<sup>a</sup>The numbers in the parentheses are the standard deviations of the rates determined in triplicate cultures

## Discussion

Viability and growth of microorganisms are largely dependent on environmental parameters such as pH and salinity (40, 41). The growth of non-halophilic neutrophilic denitrifier, *D. aromatica* strain RCB, was severely inhibited by NaCl concentrations and pH equal to or higher than 7 g L^-1^ and 8.0, respectively. Accumulation of N_2_O in substantial amounts (>1% of initially added NO_3_^-^) occurred only when strain RCB was grown at these salt and alkaline stress conditions. Transient or permanent N_2_O accumulation during denitrification under acid stress has been repeatedly observed with typical laboratory strains of canonical neutrophilic denitrifiers incubated at slightly acidic pH (~6.0) (23, 25). Both *Paracoccus denitrificans* and *Shewanella loihica* exhibited longer lag periods and slower metabolic turnover of NO_3_^-^ at pH 6.0 than at their optimal pH-neutral condition and a significant increase in N_2_O accumulation was accompanied with the growth inhibition. The observations in this study clearly show that salt and alkaline stress can also lead to elevated N_2_O production in canonical denitrifiers.

The mechanisms underpinning the increased N_2_O production in stressed *D. aromatica* strain RCB cultures were 1) alteration in the expression of the genes encoding different steps of the denitrification pathway and 2) altered kinetic balance between N_2_O-producing and consuming reactions. In previous experiments that examined denitrification at acidic pH, the transcription of *nosZ* and other denitrification genes were largely unaffected, as compared to the optimal growth conditions (22, 25). Contrastingly, in this study, the nosZ-to-(*nirS*_1_+*nirS*_2_) ratios were significantly higher at both alkaline and salt stress conditions at the time points where N_2_O accumulation was observed, suggesting that the altered NosZ-to-NirS ratio may have been one of the causes for the disrupted balance between N_2_O production and consumption. In interpreting the presented data, the *nosZ*-to-*nirS* transcription ratio of 1:1 needs not be directly translated to 1:1 activity ratio for NO_2_^-^ reduction and N_2_O reduction, as the reaction kinetics for the expressed enzymes may differ for NirS and NosZ (42). Nonetheless, the nosZ-to-(*nirS*_1_+*nirS*_2_) transcription ratios below 4.3 invariably overlapped with the incremental increases in the amounts of N_2_O. In the salt stress cultures, the recovery of nosZ-to-(*nirS*_1_+*nirS*_2_) ratios upon the delayed onset of the exponential phase to the values comparable to the level observed in exponential growth phase of the control experiment explained the temporal decline in the amounts of N_2_O in the vessels. The consistently low nosZ-to-(*nirS*_1_+*nirS*_2_) ratio during incubation of *D. aromatica* at pH 8.0 was also in agreement with the permanent N_2_O accumulation observed at this condition.

The results of the resting cell experiments indicated that the altered kinetic balance in the stressed *D. aromatica* cells lends another mechanistic explanation for elevated net N_2_O production in the pH- and salinity-stressed cells. The extent to which N_2_O accumulates in denitrification reaction is dependent on the differential between the rates of its production and consumption (13). The recent investigations on the kinetics of N_2_O consumption by the *nosZ*-harboring organisms have revealed broadly varying kinetics among the different groups of N_2_O-reducing organisms (36, 37, 43). The minimal N_2_O accumulation observed in previous observations of denitrifying isolates or enrichments suggest that the rates of N_2_O production are often surpassed by the rates of N_2_O consumption at non-acidic pH (13, 17, 24). Consistent with these previous observations and the denitrification progression curve of the *D. aromatica* strain RCB control culture (Fig. 1A), the rate of N_2_O consumption was significantly faster than the rate of N_2_O production in the chloramphenicol-treated resting cells at both high (1.0 mM NO_2_^-^ or N_2_O) and low (0.1 mM NO_2_^-^ or N_2_O) concentrations of the substrates. While the rates of N_2_O production were unaffected by the salt and alkaline stresses, the rates of N_2_O consumption were severely affected. That the decreases in N_2_O-reducing rates were much more pronounced at the low substrate concentration suggested a substantial change in the kinetics of N_2_O reduction. Such alterations in the whole-cell substrate utilization kinetics due to changes in the medium pH and salinity have been previously observed with ammonia oxidation in *Nitrosomonas europaea* and nitrite oxidation in *Nitrobacter agilis* (44, 45). The different responses to the salt and alkaline stresses on the kinetics of NO_2_^-^ and N_2_O reduction well explains the observed accumulation of N_2_O, as the rate of N_2_O production would exceed the rate of its reduction at the concentration ranges of NO_2_^-^ and N_2_O relevant to the denitrifying strain RCB cultures.

Whether the increased net N_2_O production upon exposure to the salt and alkaline stress is limited to the organisms closely related to *D. aromatica* or is widespread physiology among denitrifiers is yet unknown. Even with a conservative extrapolation confining this physiological behavior to the betaproteobacterial denitrifiers with *nirS* and clade II *nosZ* closely affiliated to those of *D. aromatica*, the findings in this study may still have significant implications to environmental N_2_O emissions. *Dechloromonas* spp. and closely affiliated organismal groups in the *Rhodocylaceae* family, e.g., *Azoarcus* spp. and *Azospira* spp., have been often identified as one of the dominant denitrifier groups in soil and freshwater environments (42, 46, 47). Recent studies using high-throughput sequencing for microbial community profiling and metagenome analyses have revealed the abundance of the *Rhodocylaceae* family of denitrifiers in estuarine environments susceptible to periodic seawater intrusions (48, 49). As estuarine environments are often enriched in reactive nitrogen due to the run-off from neighboring agricultural soils, events of seawater intrusion may result in significant N_2_O emissions from denitrification by the *Dechloromonas-like* denitrifiers exposed to salt stress. Due to climate change that will undoubtedly bring about sea level rise, larger area of estuarine deltas will be affected by seawater intrusions, and thus, the N_2_O emission mechanism examined in this study may provide significant positive feedback to global warming (27).

*Dechloromonas* spp. and closely-related organisms are also often the dominant denitrifiers in activated-sludge WWTPs, another important source of N_2_O (33, 34). Although pH of the sewers in the activated sludge tanks are relatively well maintained within the circumneutral range, temporal increases in pH to 8.0 due to, for example, shock loads of alkaline influent wastewater are not rarely observed (26). Caustic dosing often used for mitigation of sewer biofilms may also cause abrupt alkalization of activated sludge tanks (50). Such alkaline stress exerted on the *Dechloromonas*-like denitrifiers may cause an imbalance between production and consumption of N_2_O and result in substantially increased N_2_O emissions. This probable enhancement of N_2_O production upon alkaline shock may be one of the causes of highly fluctuating nature of N_2_O emissions from WWTPs (51, 52). In future research, a long-term investigation of the correlations between N_2_O emissions and fluctuating pH and expressions and activities of NirS/NirK and NosZ in an activated sludge tank may provide additional insights into N_2_O emission mechanism in activated sludge WWTPs.

## Materials and methods

### Medium and culturing conditions

A phosphate-buffered minimal salt medium containing, per liter, 0.5 g NaCl, 1.64 g Na_2_PO_4_, 1.15 g KH_2_PO_4_ and 1 mL 1000X trace metal solution was used for maintenance of the *D. aromatica* strain RCB culture and preparation of precultures for the experiments performed in this study (36, 53). The medium was boiled with continuous stream of N_2_ gas (>99.999%) to remove dissolved O_2_. Anoxic culture bottles were prepared by dispensing 96.5-mL aliquots of the cooled medium into 160-mL serum bottles using the Hungate technique (54). The serum bottles were then sealed with butyl rubber stoppers (Geo-Microbial Technologies, Inc., Ochelata, OK) and autoclaved. Sterilized anoxic stock solutions of sodium acetate, KNO_3_^-^, and NH_4_Cl were prepared separately and were added aseptically to the culture bottles to the target concentrations of 10, 5.0, and 0.5 mM, respectively. Filter-sterilized 200X vitamin stock solution was also added to the culture bottles (55). The media used for the high-pH incubation experiments (at pH 7.5 and 8.0) were prepared with different Na_2_HPO_4_:KH_2_PO_4_ ratios, although the total phosphate buffer concentration was maintained constant at 20 mM. The media with NaCl concentrations higher than 0.05% were prepared by adding appropriate volumes of 26% (w/v) NaCl solution. For each batch experiment, 0.5 mL of *D. aromatica* strain RCB preculture (pH 7.0, 0.05% NaCl) at the late exponential phase was inoculated to the prepared media bottles. All culture vessels were incubated in dark at 25°C with shaking at 160 rpm. The final pH was always within 0.2 of the initial value at any examined experimental condition.

### Cultivation of *D. aromatica* strain RCB under salt and alkaline stressed conditions

To examine whether elevated salinity or pH has any effect on N_2_O accumulation during the growth of *D. aromatica* strain RCB on denitrification, the strain RCB cells were incubated in the media with varying NaCl concentrations (0.5, 1.0, 3.0, 5.0, 7.0, 9.0, and 12 g L^-1^) and pH (7.0, 7.5, and 8.0, buffered with 20 mM phosphate). The pH of the medium was fixed to 7.0 for the salinity experiments and the NaCl concentration was fixed to 0.5 g L^-1^ for the pH experiments. The cell density (OD_600 nm_) and the amounts of N_2_O in the vessels were monitored until the end of the reaction, denoted by decrease in the cell density. All incubation experiments were performed in triplicates.

Based on the observations in these preliminary experiments, the salt stress condition with 7.0 g L^-1^ NaCl and the alkaline stress condition at pH 8.0 were selected for closer examination. The strain RCB cultures were incubated in the medium adjusted to the target NaCl concentration or pH. Upon each measurement time point, the headspace N_2_O was measured and 1 mL of the aqueous phase was withdrawn with a sterile, N_2_-flushed plastic syringe. After measuring the OD600nm, the aqueous sample was filtered using 0.2-μm-pore-size syringe filters (Advantec, Inc., Tokyo, Japan) and the filtrate was stored at −20°C until further analyses of dissolved NO_3_^-^ and NO_2_^-^. At the time points representative of different growth phases, samples were withdrawn for reverse transcription quantitative polymerase chain reaction (RT-qPCR) analyses. The extracted 400-μL cell suspensions were treated with 800 μL RNAprotect Bacteria Reagent (Qiagen, Hilden, Germany), and after centrifugation at 5,000 x g for 10 min and removal of the supernatants, the pellets were stored at −80°C until further treatment. As sample volume loss was an issue in these batch experiments, overly frequent sampling was avoided and the time intervals between sampling were maintained at approximately 48 hours. Monitoring of the culture vessels was continued until the end of the reaction, denoted by at least three consecutive measurements yielding statistically indistinguishable concentrations of the nitrogen species.

### Resting cell kinetic experiments to measure N_2_O production and consumption upon induced salt and alkalinity stress conditions

The kinetics of N_2_O production from NO_2_^-^ reduction and N_2_O consumption were examined at the salt stress (7.0 g L^-1^ NaCl) and alkaline stress (pH 8.0) conditions with the resting *D. aromatica* strain RCB cells cultivated at the control conditions. All kinetics experiments were performed in triplicate with 100-mL *D. aromatica* strain RCB cultures in 160-mL serum bottles with N_2_ headspace. The strain RCB cultures were initially incubated at the control condition with 10 mM acetate and 5 mM NO_3_^-^ as the electron donor and electron acceptor, respectively. Immediately after depletion of the initially added NO_3_^-^ and the intermediates (NO_2_^-^ and N_2_O), chloramphenicol (Sigma Aldrich, St. Louis, MO) was added to the concentration of 35 μg mL^-1^ to inhibit *de novo* protein synthesis (56). The absence of growth and linear N_2_O consumption in chloramphenicol-treated *D. aromatica* cells verified the effectiveness of chloramphenicol at the concentration tested (data not shown). The salt stress samples were prepared by adjusting the dissolved NaCl concentrations to 7.0 g L^-1^ with degassed 260 g L^-1^ NaCl brine, and the pH stress samples by adjusting the pH to 8.0 with degassed 0.5 M NaOH solution. An 1.0-mL aliquot was withdrawn from each vessel and stored at −20°C for determination of the total protein concentration.

Four sets of experiments were performed with these resting cells to determine the N_2_O production rates from NO_2_^-^ reduction and N_2_O consumption rates at two different initial dissolved concentrations. For determination of the N_2_O production rates, NaNO_2_ was added to the resting cells to the concentration of 0.1 mM or 1.0 mM, and 6.0 mL of the N_2_ headspace was replaced with 99.9% acetylene gas (Taekyung Eco Co., Ansan, Korea) to inhibit N_2_O reduction (57). The culture vessels for measurement of N_2_O consumption rates were prepared by adding autoclaved 99.999% N_2_O and 10% v/v N_2_O in N_2_ to the headspace to yield the target aqueous N_2_O concentration of 1.0 mM and 0.1 mM, respectively. The increase or decrease in N_2_O concentrations in these vessels were monitored until the changes in the concentrations were no longer linear, and the reaction rates were determined from linear regression of the N_2_O production or consumption curves. These N_2_O production and consumption rates were normalized with total protein concentration determined with the Bradford protein assay using Bio-Rad Protein Assay Kit II (Hercules, CA) according to the instruction provided by the manufacturer.

### Analytical procedures

Aqueous concentrations of NO_3_-N and NO_2_^-^-N were determined with the spectrophotometric detection method using the Griess reagents as previously described (58). Vanadium chloride (VCl3) was used as the reducing agent to reduce NO_3_^-^ to NO_2_^-^. The absorbance at 540 nm was measured using a Sunrise absorbance microplate reader (Tecan, Männedorf, Switzerland). The headspace N_2_O concentrations were monitored using a HP6890 series gas chromatograph equipped with an HP-PLOT/Q column and electron capture detector (Agilent, Palo Alto, CA). For each measurement, 200 μL of the headspace gas was withdrawn and manually injected into the gas chromatograph with a 1700 series gas-tight syringe (Hamilton Company, Reno, NV). The total amount of N_2_O in a culture vessel was calculated from the headspace concentration with the Henry’s constant adjusted for the salinity of the medium, as previously described (59–62). The cell density was determined by measuring the absorbance at 600 nm with a spectrophotometer (Thermo Scientific, Waltham, MA, United States).

### Reverse transcription quantitative PCR (RT-qPCR) analyses

The transcription profiles of the genes encoding the enzymes involved in denitrification at different incubation conditions were analyzed using RT-qPCR analyses performed using the previously described protocol (Table 3) (63). The RNAprotect-treated samples stored at −80°C were thawed on ice and one-microliter solution with 10^10^ copies of luciferase control mRNA (Promega, Madison, WI) was added to each sample tube. Total RNA was extracted according to the protocol provided with the RNeasy Mini Kit (Qiagen). The total RNA collected in the column was eluted twice with 30 μL RNase-free water, and the resulting eluent was treated with RNase-free DNAse Set (Qiagen) and subsequently purified with RNeasy MinElute Kit (Qiagen). Of the 20 μL eluent, 10 μL was subjected to reverse transcription using Superscript^™^ III Reverse Transcriptase (Invitrogen, Carlsbad, CA) as previously described (63). The rest of the eluent was stored at −20 °C and later used for confirmation of absence of DNA in the purified RNA samples. After reverse transcription, 1 μL of RNase H (Invitrogen) was added to each reaction mixture, which was then incubated at 37 °C for 20 minutes. The cDNA solution was diluted 5-fold with nuclease-free water and stored at −20°C.

**Table 3.**
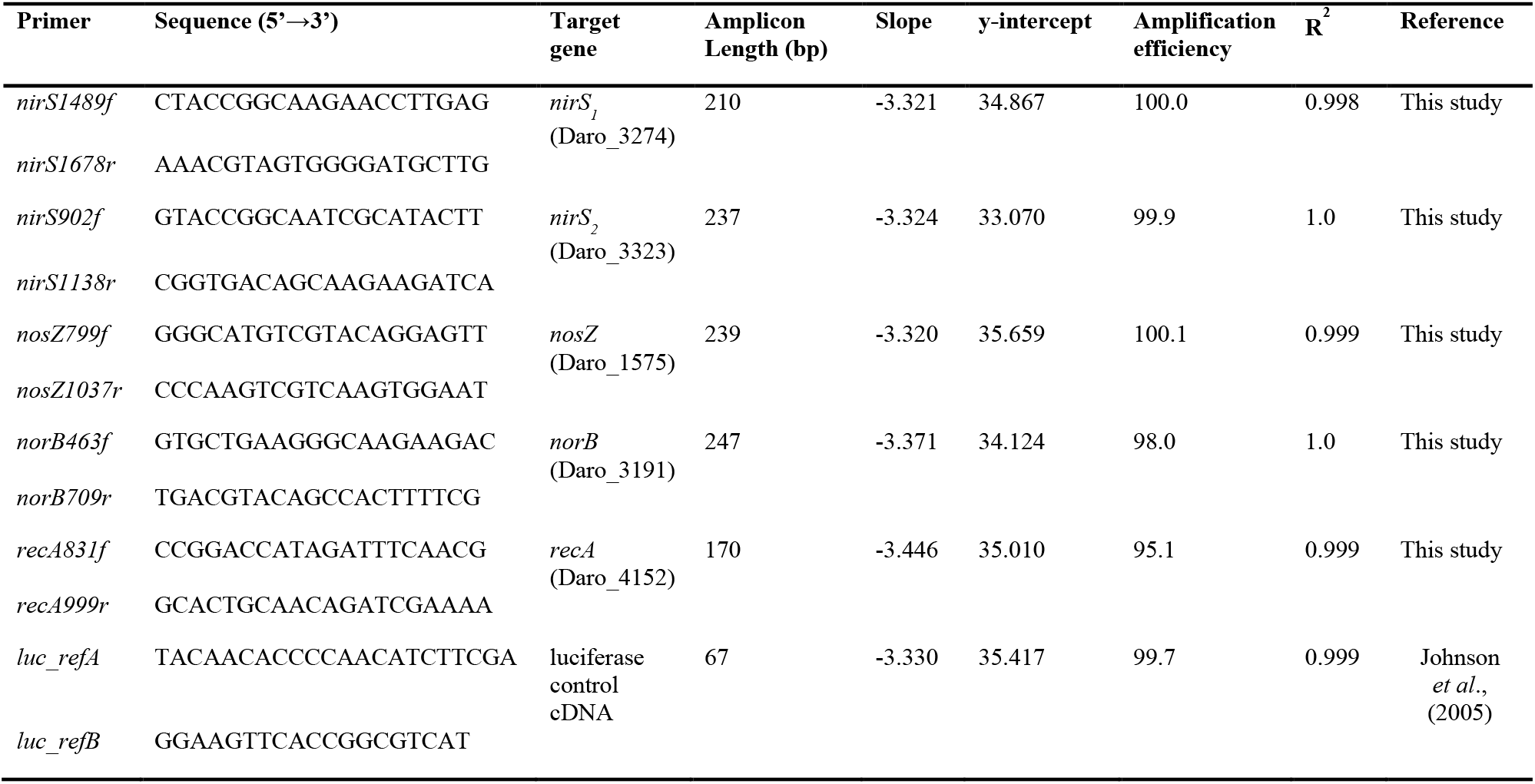
The primer sets used for the RT-qPCR assays.

The primer sets used for the RT-qPCR analyses were designed *de novo* from the genome sequence of *D. aromatica* strain RCB available at the NCBI genome database (accession number: CP000089.1) using Primer3 software (64). The primers targeting *nirS*_1_ (Daro_3274), *nirS*_2_ (Daro_3323), *norB* (Daro_3191), *nosZ* (Daro_1575), and *recA* (Daro_4152) were designed and the calibration curves were constructed with the dilution series of PCR^™^2.1 plasmids (Invitrogen) with amplicons inserted (Table 3). Amplification was performed with QuantStudio^™^ 3 Real-Time PCR System (Thermo Fisher Scientific) using SYBR Green detection chemistry. The remaining eluent stored after the DNase treatment step was analyzed along with the cDNA samples, to confirm that DNA was completely removed by the DNase treatment. The copy numbers of *luc* genes in the cDNA samples were quantified to ensure that the RNA recovery rates were acceptable (>10%). The transcript copy numbers of *nirS*_1_, *nirS*_2_, *norB*, and *nosZ* were normalized with the transcript copy number of *recA* (65). The samples withdrawn from triplicate cultures were independently treated through the RNA extraction and reverse transcription steps, serving as the biological replicates of the RT-qPCR analyses. Each qPCR reaction was performed in duplicate and the average CT value of the technical replicates was taken.

### Statistical analyses

The statistical analyses in this study was performed with the R statistical software package (version 3.5.0). The statistical comparisons between two sets of data were performed with t-tests and one-way analysis of variance (ANOVA) was used for analyses of three or more sets of data. Unless otherwise mentioned, triplicate experiments were performed and the average values were presented with the standard deviations of the triplicate samples as errors.

## Acknowledgements

This project is supported by the “R&D Center for reduction of Non-CO_2_ Greenhouse gases (2017002420002)” funded by Korea Ministry of Environment (MOE) as “Global Top Environment R&D Program”. The authors report no conflict of interest.

